# SNVPhyl: A Single Nucleotide Variant Phylogenomics pipeline for microbial genomic epidemiology

**DOI:** 10.1101/092940

**Authors:** Aaron Petkau, Philip Mabon, Cameron Sieffert, Natalie Knox, Jennifer Cabral, Mariam Iskander, Mark Iskander, Kelly Weedmark, Rahat Zaheer, Lee S. Katz, Celine Nadon, Aleisha Reimer, Eduardo Taboada, Robert G. Beiko, William Hsiao, Fiona Brinkman, Morag Graham, the IRIDA Consortium, Gary Van Domselaar

## Abstract

**Motivation:** The recent widespread application of whole-genome sequencing (WGS) for microbial disease investigations has spurred the development of new bioinformatics tools, including a notable proliferation of phylogenomics pipelines designed for infectious disease surveillance and outbreak investigation. Transitioning the use of WGS data out of the research lab and into the front lines of surveillance and outbreak response requires user-friendly, reproducible, and scalable pipelines that have been well validated.

**Results:** SNVPhyl (Single Nucleotide Variant Phylogenomics) is a bioinformatics pipeline for identifying high-quality SNVs and constructing a whole genome phylogeny from a collection of WGS reads and a reference genome. Individual pipeline components are integrated into the Galaxy bioinformatics framework, enabling data analysis in a user-friendly, reproducible, and scalable environment. We show that SNVPhyl can detect SNVs with high sensitivity and specificity and identify and remove regions of high SNV density (indicative of recombination). SNVPhyl is able to correctly distinguish outbreak from non-outbreak isolates across a range of variant-calling settings, sequencing-coverage thresholds, or in the presence of contamination.

**Availability:** SNVPhyl is available as a Galaxy workflow, Docker and virtual machine images, and a Unix-based command-line application. SNVPhyl is released under the Apache 2.0 license and available at http://snvphyl.readthedocs.io/ or at https://github.com/phac-nml/snvphyl-galaxy.

## Introduction

The high-efficiency and cost-effectiveness of whole-genome sequencing (WGS) using next-generation sequencing (NGS) technologies is transforming the biomedical landscape. Entire microbial genomes can be rapidly sequenced and subsequently queried with nucleotide-level resolution, an exciting new ability that far outstrips other traditional microbial typing methods. This powerful new ability has the potential to advance many fields, including in particular the field of infectious disease genomic epidemiology. A number of landmark studies have demonstrated the power of WGS for molecular epidemiology. One notable study is the investigation into the 2010 Haiti cholera outbreak (1–3), where WGS and epidemiological data was used in support of the hypothesis that cholera was introduced to Haiti from UN peacekeepers originally infected in Nepal. WGS has supported the investigation of outbreaks of organisms as diverse as *Mycobacterium tuberculosis* (4, 5), *Escherichia coli* (6), and *Legionella pneumophila* (7). These high-profile successes have motivated public health institutions and food regulatory agencies to incorporate WGS into their routine microbial infectious disease surveillance and outbreak investigation activities. The GenomeTrakr network used by the Centers for Disease Control (CDC) and the Food and Drug Administration (FDA) agencies in the United States (8), PulseNet International (http://www.cdc.gov/pulsenet/next-generation.html), Statens Serum Institut in Denmark (9), and Public Health England (10) are leading the charge in this area, and have incorporated a variety of analytical approaches to integrate WGS into their infectious disease surveillance activities. Two approaches in particular have emerged as feasible methods for bacterial genomic epidemiology: gene-by-gene methods, which extend the idea of multilocus sequence typing (MLST) to encompass a given organism’s entire genome (whole-genome MLST, wgMLST) or core genome (core genome MLST, cgMLST) (11, 12); and single nucleotide variant (SNV)-based methods, which identify variants by comparing a population of target genomes against a reference (13, 14). Gene-by gene methods are promising as they are more amenable to assigning consistent sequence types using standardized MLST schemas, but these schemas must be developed and validated for each organism. SNV-based methods are popular as they do not require development of MLST schemas, but the variability in SNV-identification methods and reference genome selection means they do not yet produce standard sequence types useful for global communication of circulating infectious disease (12, 14). Where applicable, these two methods are often combined (15).

A growing number of SNV-based pipelines have been developed (Table 1) and are distributed in the form of web services (16), command-line software (17), or both (14). Web services provide a user-friendly method of running large-scale analyses but require the uploading of sequence reads and rely on third-party computing infrastructure, which may be inadequate for the analysis of typically large datasets or due to data privacy concerns. Locally installed pipelines avoid the transfer of large datasets to third-party websites, offer greater control over the execution environment for reproducibility, and allow for the incorporation into pre-existing bioinformatics analysis environments. However, locally installed pipelines may require considerable expertise to operate and can have substantial computing requirements. Additionally, for many SNV-based pipelines, recombination detection and removal may require pre-analysis to identify phage and genomic islands on the reference genome, or post-analysis with computationally intensive recombination-detection software such as Gubbins (18) or ClonalFrameML (19) to identify and mask possible recombinant regions. While a large choice of pipelines is available, a systematic comparison of popular SNV pipelines has demonstrated that they generate highly concordant phylogenetic trees but with variation in the particular SNVs identified (20). However, variation in the installation procedures and execution environments of these pipelines proves challenging for integration into a larger bioinformatics analysis system.

**Table 1.** A comparison of whole-genome phylogenetic software.

| Name | Input<sup>1</sup> | Parallel Computing<sup>2</sup> | Distribution<sup>3</sup> | Interface<sup>4</sup> | Reference |
| --- | --- | --- | --- | --- | --- |
| CFSAN SNP Pipeline | sr | mn,mt | local | cl | (17) |
| CSIPhylogeny | sr,ag | n/a | web | gui | (16) |
| kSNP | sr,ag | mt | local | cl | (43) |
| Lyve-SET | sr,ag* | mn,mt | local | cl | https://github.com/lskatz/lyve-SET |
| NASP | sr,ag | mn,mt | local | cl | (20) |
| Parsnp | ag | mt | local | cl | (44) |
| PhaME | sr,ag | mt | local | cl | (45) |
| REALPHY | sr,ag | mt | web,local | gui,cl | (14) |
| Snippy | sr,ag* | mt | local | cl | https://github.com/tseemann/snippy |
| SNVPhyl | sr | mn,mt | local | gui,cl | http://snvphyl.readthedocs.io/ |

<sup>1</sup>sr = sequence reads, ag = assembled genome, ag\* = assembled genome supported by generating simulated reads
<sup>2</sup>mn (multi-node) = provides capability to execute across multiple compute nodes, mt (multi-thread) = provides multi-threading capability, n/a = not applicable (not locally installable)
<sup>3</sup>local = locally distributed and installable software, web = software provided as a web service
<sup>4</sup>cl = command-line interface, gui = graphical user interface

Galaxy (21) is a web-based biological data analysis platform that can be accessed through a publicly available website, a locally installed instance linked to a high-performance compute cluster, or a cloud-based environment. Galaxy provides a user-friendly web interface for the construction of data analysis workflows using a mixture of built-in or community developed bioinformatics tools. Additionally, Galaxy provides an API for automated workflow execution or other automations via external software. These features have encouraged some software developers to integrate Galaxy within larger data analysis systems. Examples of such analysis systems include IRIDA (http://irida.ca), the Refinery Platform (http://www.refinery-platform.org/), and the Genomics Virtual Laboratory (22).

The SNVPhyl pipeline provides a reference-based SNV discovery and phylogenomic tree-building pipeline along with ancillary tools integrated within the Galaxy framework. SNVPhyl can quickly analyze many genomes, identify variants and generate a maximum likelihood phylogeny, an all-against-all SNV distance matrix, as well as additional quality information to help guide interpretation of the results. The pipeline has been under continuous development and refinement at Canada's National Microbiology Laboratory since 2010; it is currently being used for outbreak investigations and will be part of the validated suite of tools used by PulseNet Canada for routine foodborne disease surveillance activities. Here, we describe the overall operation of SNVPhyl, survey its advanced features such as repeat and recombination masking, and demonstrate its SNV-calling and phylogenomic tree building accuracy using simulated and real-world datasets.

## Methods

### SNVPhyl pipeline

The SNVPhyl pipeline (Figure 1) consists of a set of pre-existing and custom-developed bioinformatics tools for reference mapping, variant discovery, and phylogeny construction from identified SNVs. Each stage of the pipeline is implemented as a separate Galaxy tool and the stages are joined together to construct the SNVPhyl workflow. Distribution of the dependency tools for SNVPhyl is managed through the Galaxy Toolshed (23). Scheduling of each tool is managed by Galaxy, which provides support for execution on a single machine, high-performance computing environments utilizing most major scheduling engines (e.g., Slurm, TORQUE, Open Grid Engine), or cloud-based environments.

**Figure 1.**
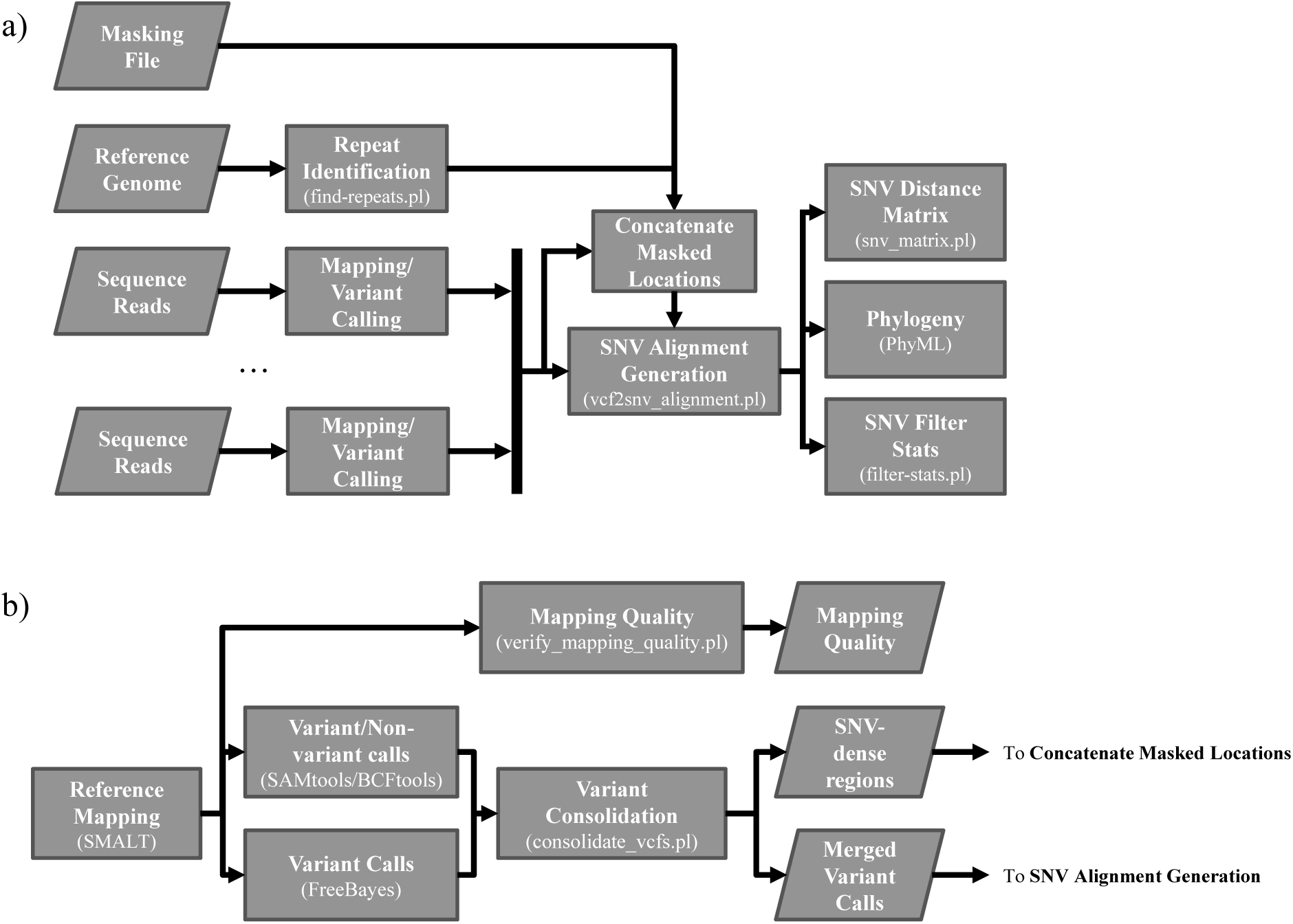
**a)** Overview of the SNVPhyl pipeline. Input to the pipeline is provided as a reference genome, set of sequence reads for each isolate, and an optional list of positions to mask from the final results. Repeat regions are identified on the reference genome and reference mapping followed by variant calling is performed on the sequence reads. The resulting files are compiled together to construct a SNV alignment and list of identified SNVs, which are further processed to construct a SNV distance matrix, maximum likelihood phylogeny, and a summary of the identified SNVs. Individual software or scripts are given in the parenthesis below each stage. **b)** An overview of the “Mapping/Variant Calling” stage of SNVPhyl. Variants are called using two separate software packages and compiled together in the “Variant Consolidation” stage. As output, a list of the validated variant calls, regions with high-density SNVs, as well as quality information on the mean mapping coverage are produced and sent to further stages.

### Input

SNVPhyl requires as input a set of microbial WGS datasets, a reference genome, and an optional masking file defining regions on the reference genome to exclude from the analysis. The sequencing data consists of either single-end or paired-end reads. The reference genome consists of a draft or finished genome, chosen typically to have high similarity with the collection of genome sequences under analysis. The masking file stores the sequence identifier of the reference genome and the coordinates for any regions where SNVs should be excluded from analysis.

### Architecture

Execution of SNVPhyl begins with the “Repeat Identification” stage. This stage identifies internal repeat regions on the reference genome using MUMMer (v3.23) (24) and generates a masking file containing the locations of repetitive regions to exclude from analysis. This file is concatenated to the user-supplied masking file, if defined, and used in later analysis stages.

The “Mapping/Variant Calling” stage (detailed in Figure 1.b) aligns the supplied reads to the reference genome using the appropriate mapping mode (paired-end or single-end). Reference mapping is performed using SMALT (v.0.7.5) (http://www.sanger.ac.uk/science/tools/smalt-0), which outputs a read pileup. In the “Mapping Quality” stage, SNVPhyl evaluates each pileup for the mean coverage across a user-defined proportion of the reference genome (e.g., 10X coverage across at least 80% of the genome). Any sequenced genomes that do not meet the minimum mean coverage threshold are flagged for further assessment.

The variant calling stages of SNVPhyl use two independent variant callers, FreeBayes (version 0.9.20) (25), and the SAMtools and BCFtools packages (26, 27). FreeBayes is run using the haploid variant calling mode and the resulting variants are filtered to remove insertions/deletions and split complex variant calls. SAMtools and BCFtools are run independently of FreeBayes and are used to confirm the FreeBayes variant calls and generate base calls for non-variant positions.

The “Variant Consolidation” stage combines both sets of variant and non-variant calls into a merged file, flagging mismatches between variant callers. Base calls below the defined minimum read coverage are identified and flagged. The merged base calls are scanned for positions that do not pass the minimum SNV abundance ratio (ratio of reads supporting the SNV with respect to the depth of coverage at a site) and minimum mean mapping quality. These base calls are removed from the merged base calls file. The remaining base calls that pass all these criteria are defined as either a high quality SNV (hqSNV) or a high quality non-variant base call. The hqSNVs are optionally scanned to identify high density SNV regions. These regions are identified by passing a sliding window of a given size along the genome and counting the number of SNVs within the window that exceed a given SNV density threshold. The high-density SNV regions are recorded in a tab-delimited file and used to mask potential recombinant regions.

The “SNV Alignment Generation” stage examines the merged base calls to generate a table of identified variants and an alignment of hqSNVs and high-quality non-variant bases. The hqSNVs are evaluated and assigned a status using the base calls at the same reference genome position for every isolate. A status of “valid” is assigned when the base calls from all isolates in the same position pass the minimum criteria (hqSNVs or high-quality non-variants). These base calls are incorporated into the SNV alignment used for phylogeny generation. A status of “filtered-coverage” is assigned when one or more isolates fail the minimum coverage threshold and the failed isolates’ base calls are annotated as ‘-‘ (indicating no nucleotide or a gap). A status of “filtered-mpileup” is assigned when one or more isolates have conflicting base calls between FreeBayes and SAMtools/BCFtools and the conflicting isolates’ base calls are annotated as ‘N’ (indicating any nucleotide nonspecifically). A status of “filtered-invalid” is assigned when the identified hqSNV overlaps one of the masked locations. The hqSNVs, base calls, and assigned status are recorded in the SNV table and saved for later inspection. The SNV table can be used to re-generate the downstream SNV alignment and phylogenetic tree without re-running the computationally intensive reference mapping and variant calling steps.

### Output

The final phylogeny is generated using the SNV alignment consisting of hqSNVs with a “valid” status. This alignment is run through PhyML (28) with the GTR+ *γ* model as default and tree support values estimated using PhyML’s approximate likelihood ratio test (29). The SNV alignment is also used to generate an all-against-all SNV distance matrix. This matrix lists the pair-wise distances between every isolate, using only the “valid” hqSNVs.

Additional files are provided to assist in evaluating the quality of the SNVPhyl analysis. The “SNV Filter Stats” stage summarizes the quality and counts of the identified SNVs. The “SNV Alignment Generation” stage summarizes the proportion of the reference genome passing all the necessary filters for every isolate—the (non-masked) core genome—as well as the portion of the genome failing any filter or excluded by the masking file.

### Simulated data

We evaluated SNVPhyl’s sensitivity and specificity for SNV identification using simulated mutations derived from a reference genome. The closed and finished *E. coli* str. Sakai (NC_002695) along with the two plasmids (NC_002128 and NC_002127) was chosen as the reference genome (combined length of 5,594,477 bp). We constructed a variant genome by randomly mutating 10,000 base locations on the reference genome. We repeated the procedure, using the same 10,000 base locations but different mutations, to generate a total of three variant genomes. We included the unmodified reference genome in the test set to serve as a positive control. The simulated variants for each genome were recorded in a table for later comparisons. The constructed genomes were run through art_illumina (version ChocolateCherryCake) (30) to generate paired-end reads with 2x250 bp length and 30X mean coverage. The resultant reads along with the reference genome were run through SNVPhyl with repeat masking enabled but with no SNV density filtering.

The SNV table produced by SNVPhyl was compared to the table of simulated variants to determine their sensitivity and specificity. We define a true positive (TP) as a matching row in both variant tables where both the position as well as base calls for each simulated genome is identical. A variant detected by SNVPhyl not matching the criteria for a true positive is a false positive (FP). A true negative (TN) is defined as all non-variant positions that were excluded by SNVPhyl. A false negative (FN) is defined as a row in the simulated variant table where either the position or a base call did not match any corresponding entry in the table of detected variants by SNVPhyl. Using these definitions, sensitivity is calculated as TP/(TP + FN) while specificity is calculated as TN/(TN + FP).

### SNV density filtering evaluation

We evaluated SNVPhyl’s ability to mask recombination by comparing the resultant phylogenetic trees and identified SNVs to those detected and removed by the recombination detection software package Gubbins (18). Our test data consisted of 11 *Streptococcus pneumoniae* genomes along with the reference genome ATCC 700669 (FM211187) that had previously been published (31) and made available as sequence reads on NCBI (Table S1) and as a whole-genome alignment (the PMEN1 dataset from https://sanger-pathogens.github.io/gubbins/). We downloaded this alignment, appended the reference genome, and processed the resulting file through Gubbins to identify and mask recombinant SNVs. The identified SNVs were filtered to remove gaps and masked recombination (‘-’ and ‘N’ characters) and the resulting SNVs we defined as the “truth” set used to generate the true/false positive/negative values—defined as for the “Simulated data” section. These Gubbins-identified SNVs were also used to construct a phylogenetic tree with PhyML and compared with SNVPhyl’s phylogenetic trees numerically using K tree scores (32) and visually using phytools (33). K tree scores allow for similarity comparisons of many phylogenetic trees against a single reference tree. Each tree is re-scaled by a factor, K, based on the reference tree size and a score is produced taking into account differences in both topology and branch lengths. Comparing the scores of all trees provides a measure of similarity to the reference tree, with more similar trees producing a score closer to 0.

We downloaded sequence reads for the test dataset from NCBI, identifying and combining multiple sequencing runs for each strain to a single set of sequence reads with the help of SRAdb (34). Using the combined sequence reads we ran SNVPhyl under a number of scenarios. For each scenario we compared the SNVs and phylogenetic trees to the “truth” dataset described above. In the first run, we performed no SNV density filtering. For all subsequent runs we adjusted the density-filtering parameters to remove SNVs occurring at a density of 2 or more within a moving window of 20, 100, 500, 1000, and 2000 bp. We evaluated an additional scenario using a combination of SNVPhyl and Gubbins for recombination masking. We ran SNVPhyl with no SNV density filtering and incorporated the identified
variants into the reference genome to generate a whole genome alignment. The whole-genome alignment was processed with Gubbins to identify non-recombinant SNVs and to construct a phylogenetic tree.

### Parameter optimization

We evaluated SNVPhyl’s parameter settings and resulting accuracy at differentiating outbreak isolates using a set of 59 sequenced and published *Salmonella enterica* serovar Heidelberg genomes (35), which were previously deposited in the NCBI Sequence Read Archive (Table S2). We chose this dataset as it contained sequence data for strains from several unrelated outbreaks—referred to as “outbreak 1”, “outbreak 2”, and “outbreak 3”—along with additional background strains, allowing us to evaluate SNVPhyl’s ability to differentiate the outbreak strains under different scenarios. Sequence read data was subsampled with seqtk (https://github.com/lh3/seqtk) such that the genome with the least amount of sequence data, SH12-006, was set to 30X mean coverage (calculated as: mean coverage = count of base pairs in all reads / length of reference genome). Other genomes were subsampled to maintain their relative proportion of mean read coverage to SH12-006. *Salmonella* Heidelberg str. SL476 (NC_011083) was selected as the reference genome. We optimized the SNVPhyl parameters for this dataset according to the following four scenarios: 1) adjusting the minimum base coverage parameter used to call a variant while keeping the number of reads in the dataset fixed; 2) subsampling the reads of a single WGS sample at different mean coverage levels while keeping the minimum base coverage parameter fixed; 3) adjusting the minimum SNV abundance ratio for calling a variant; and 4) adjusting the amount of contamination in the dataset to determine its effect on variant calling accuracy.

In the first scenario we ran the SNVPhyl pipeline using the default parameters except for the minimum base coverage, which was adjusted to 5X, 10X, 15X, and 20X. In the second scenario we kept the minimum base coverage parameter fixed at 10X, while one of the samples (SH13-001) was subsampled to mean sequencing coverages of 30X, 20X, 15X, and 10X. In the third scenario the minimum SNV abundance ratio was adjusted to 0.25, 0.5, 0.75, and 0.9. In the fourth scenario, a sample from “outbreak 2” (SH13-001 with mean coverage 71X) was chosen as a candidate for simulating contamination. A sample from the unrelated “outbreak 1” (SH12-001) was selected as the source of contaminant reads. The reads were subsampled and combined such that SH13-001 (“outbreak 2”) remained at 71X mean coverage but was contaminated with reads from SH12-001 (“outbreak 1”) at 5%, 10%, 20%, and 30%. All samples were run through SNVPhyl for each of these contamination ratios.

The phylogenetic trees produced by SNVPhyl were evaluated for concordance with the outbreak epidemiological data using the following criteria: 1) all outbreak isolates group monophyletically, and 2) the SNV distance between any two isolates within an outbreak clade is less than 5 SNVs, a number identified in the previous study (35) as the maximum SNV distance between epidemiologically related samples within these particular outbreaks. Both conditions were tested using the APE package within R (36).

## Results

### Validation against simulated data

We measured SNVPhyl‘s sensitivity and specificity by introducing random mutations along the *E. coli* Sakai reference genome and compared these mutations with those detected by SNVPhyl (Table 2). Of the 10,000 mutated positions introduced, SNVPhyl reported 9,116 true positives and 0 false positives resulting in a sensitivity and specificity of 0.91 and 1.0.

**Table 2.** SNV simulation results.

| Comparison | Variant columns simulated | Non-variant columns | True positives | False positives | True negatives | False negatives | Specificity | Sensitivity |
| --- | --- | --- | --- | --- | --- | --- | --- | --- |
| Valid SNVs<sup>1</sup> | 10000 | 5584477 | 9116 | 0 | 5575361 | 884 | 1.0 | 0.91 |
| All SNVs<sup>2</sup> | 10000 | 5584477 | 9573 | 51 | 5574853 | 427 | 1.0 | 0.96 |

<sup>1</sup>Valid SNVs: The number of SNV-containing sites detected that passed all thresholds to be considered high quality for every isolate.
<sup>2</sup>All SNVs: All the SNV-containing sites identified by SNVPhyl, including those where at least one isolate did not have a high-quality base call or sites that were masked by the pipeline.

Positions on the reference genome that contain a low-quality base call, or exist in repetitive regions are excluded from downstream analysis by SNVPhyl. However, lower-quality variant-containing sites along with variants in repetitive regions are saved by SNVPhyl in the variant table with a “filtered” status. Comparing the combination of low-quality and high-quality variant sites detected by SNVPhyl, we found 9,573 true positives and 51 false positives resulting in a sensitivity and specificity of 0.96 and 1.0 (after rounding). Of the 51 false positives, 48 were considered as false positives due to insufficient read coverage in one of the samples to call a high quality variant, thus resulting in a call of a gap (‘-‘) as opposed to the true base call. Only 3 of the false positives were a result of miscalled bases with sufficient read coverage, and these occurred in repetitive regions of the genome.

### SNV density filtering evaluation

We compared SNVPhyl’s density filtering against the Gubbins software for detection and removal of recombination in a collection of WGS reads from 11 *Streptococcus pneumoniae* genomes along with the reference genome ATCC 700669 (Table 3, Figure S1). With no SNV density filtering SNVPhyl properly identified 142 SNV-containing sites (true positives) but included 2,159 additional SNV sites (false positives). These false positives skew the resulting phylogenetic tree by increasing the length of one of the branches. The phylogenetic tree is compared with the tree produced with Gubbins, resulting in a K tree score of 0.419.

**Table 3.**
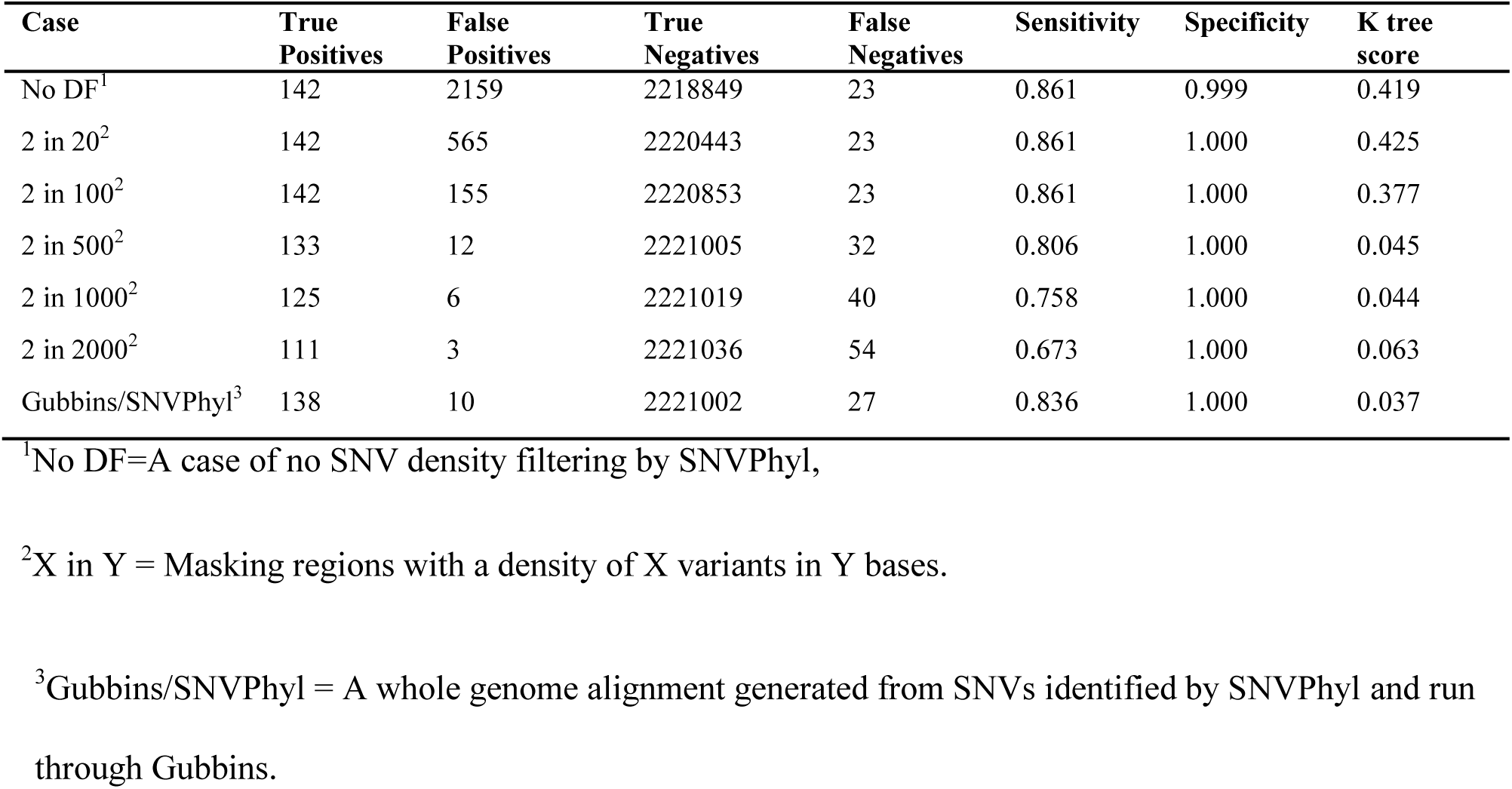
A comparison of the SNVPhyl variant density filtering algorithm to the Gubbins system for recombination detection.

| Case | True Positives | False Positives | True Negatives | False Negatives | Sensitivity | Specificity | K tree score |
| --- | --- | --- | --- | --- | --- | --- | --- |
| No DF<sup>1</sup> | 142 | 2159 | 2218849 | 23 | 0.861 | 0.999 | 0.419 |
| 2 in 20<sup>2</sup> | 142 | 565 | 2220443 | 23 | 0.861 | 1.000 | 0.425 |
| 2 in 100<sup>2</sup> | 142 | 155 | 2220853 | 23 | 0.861 | 1.000 | 0.377 |
| 2 in 500<sup>2</sup> | 133 | 12 | 2221005 | 32 | 0.806 | 1.000 | 0.045 |
| 2 in 1000<sup>2</sup> | 125 | 6 | 2221019 | 40 | 0.758 | 1.000 | 0.044 |
| 2 in 2000<sup>2</sup> | 111 | 3 | 2221036 | 54 | 0.673 | 1.000 | 0.063 |
| Gubbins/SNVPhyl<sup>3</sup> | 138 | 10 | 2221002 | 27 | 0.836 | 1.000 | 0.037 |

<sup>1</sup>No DF=A case of no SNV density filtering by SNVPhyl,
<sup>2</sup>X in Y = Masking regions with a density of X variants in Y bases.
<sup>3</sup>Gubbins/SNVPhyl = A whole genome alignment generated from SNVs identified by SNVPhyl and run through Gubbins.

We reanalyzed the dataset with high-density SNV masking enabled, using a range of variant density cutoffs. We found the density-filtering criteria of 2 SNVs in a 500 bp window and 2 SNVs in a 1000 bp window performed near-equally in producing a phylogenetic tree resembling the tree produce by Gubbins based on the K tree scores of 0.045 and 0.044, both much lower than the score of 0.419 for no SNV density filtering. With these filtering criteria, SNVPhyl identified 133 true positives and 12 false positives (for 2 SNVs in 500 bp) and 125 true positives and 6 false positives (for 2 SNVs in 1000 bp).

We also investigated the effect of generating a whole-genome alignment—by incorporating SNVPhyl-identified variants without SNV density filtering into the reference genome—for a more thorough analysis with the recombination-detection software Gubbins. We were able to identify 138 true positives in the alignment at the expense of 10 false positives and a K tree score of 0.037, a result closely matching the use SNVPhyl’s density filtering criteria.

### Parameter optimization

We evaluated SNVPhyl’s capability to differentiate between epidemiologically related and unrelated samples using a WGS dataset consisting of 59 *Salmonella enterica* serovar Heidelberg genomes from three unrelated outbreaks. We ran SNVPhyl with this data under a number of scenarios: 1) varying the minimum base coverage required by SNVPhyl to call a variant, 2) subsampling the reads of an individual bacterial sample, 3) varying the minimum SNV abundance ratio, and 4) testing the ability to generate accurate phylogenetic trees in the presence of contamination. We tested the SNVPhyl results for phylogenetic concordance to epidemiological data (Table 4, Figure S2).

**Table 4.** A comparison of the performance of SNVPhyl across a range of parameters and analysis scenarios.

| # | Scenario | Parameters/Conditions | hqSNVs | % Core | Differentiated Outbreaks |
| --- | --- | --- | --- | --- | --- |
| 1 | **Minimum Coverage** | 5X | 317 | 95 | Yes |
|  |  | 10X | 301 | 92 | Yes |
|  |  | 15X | 262 | 81 | Yes |
|  |  | 20X | 165 | 54 | No |
| 2 | **Subsample Coverage Level** | 10X<sup>1</sup> | 155 | 47 | No |
|  |  | 15X<sup>1</sup> | 242 | 76 | Yes |
|  |  | 20X<sup>1</sup> | 276 | 88 | Yes |
|  |  | 30X<sup>1</sup> | 299 | 92 | Yes |
| 3 | **SNV Abundance Ratio** | 0.25 | 351 | 92 | No |
|  |  | 0.5 | 307 | 92 | No |
|  |  | 0.75 | 301 | 92 | Yes |
|  |  | 0.9 | 291 | 92 | Yes |
| 4 | **Contamination** | 5% | 298 | 92 | Yes |
|  |  | 10% | 292 | 92 | Yes |
|  |  | 20% | 260 | 92 | No |
|  |  | 30% | 231 | 92 | No |

<sup>1</sup>These represent the mean coverage of one sample after subsampling reads and not the minimum coverage parameter of SNVPhyl (which is fixed at 10X).

For the first scenario we found that as the minimum coverage threshold was increased, the percent of the reference genome identified as part of the core genome and number of SNV-containing sites was reduced (from 95% core and 317 SNVs to 54% core and 165 SNVs). At 15X minimum coverage (81% core and 262 SNVs) and lower all three outbreaks grouped into monophyletic clades. Failure occurred at a minimum coverage of 20X (54% core and 165 SNVs), where the outbreak 2 isolates failed to constitute a separate clade.

For the second scenario, one of the samples was subsampled to reduce the mean coverage relative to all other samples while keeping the minimum coverage parameter of 10X in SNVPhyl fixed. At a mean coverage of 15X (with 242 SNVs identified and 76% core) SNVPhyl grouped all three outbreaks into monophyletic clades. However, at a lower mean coverage of 10X (155 SNVs and 47% core) SNVPhyl failed to group one of the outbreaks into a monophyletic clade. Similar to the first scenario, the percentage of the reference genome considered core as well as the number of SNVs identified was reduced as the mean coverage of one of the samples was lowered.

For the third scenario, the SNV abundance ratio—defining the ratio of SNV-supporting bases needed to identify a variant as high-quality—was adjusted incrementally. Each set of outbreak isolates grouped into a clade with a maximum SNV distance less than 5 SNVs above a ratio of 0.5. At a ratio of 0.5 the maximum SNV distance within outbreak 2 was exactly 5 SNVs while for a ratio of 0.25 the maximum SNV distance in outbreak 2 was 44 SNVs. The percentage of the reference genome identified as part of the core genome remained the same at 92%.

For the fourth scenario, we examined the robustness of SNVPhyl to cross-contamination of closely related samples. Current methods of contamination detection often focus on taxonomic classification of genomic content (37). However, contamination by closely related isolates can go undetected. We simulated contamination for an isolate in outbreak 2 by an isolate in outbreak 1. We found that SNVPhyl was able to accurately differentiate all three outbreaks with up to 10% read contamination; however the number of SNVs dropped from 298 SNVs at 5% contamination, to 260 SNVs at 20% contamination, where the failure was due to removal of the majority of unique SNVs that differentiated outbreak 1 from the background isolates.

## Discussion

The availability of WGS data from microbial genomes represents a tremendous opportunity for infectious disease surveillance and outbreak response. Emerging analytical methods, such as gene-by-gene or SNV-based, require that bioinformatics pipelines be designed with usability by non-bioinformaticians in mind and which can be easily incorporated into existing systems. An overview of current phylogenomic methods appears in (38) and a comparison of SNVPhyl's design with that of other popular pipelines appears in Table 1 (a detailed investigation comparing the performance of SNVPhyl with other pipelines is the subject of a forthcoming manuscript). We designed SNVPhyl to be both flexible and scalable in its usage in order to meet the needs and abilities of most labs. SNVPhyl gains much of this flexibility through its implementation as a Galaxy workflow, which enables execution in environments from single machines to high-scale computer clusters, from third-party web-based environments to local installations. Galaxy provides a user-friendly interface but also provides an API, which is used to implement a command-line interface for SNVPhyl. The SNVPhyl pipeline is also integrated within the IRIDA platform (http://irida.ca), which provides an integrated “push-button” system for genomic epidemiology. However, implementing SNVPhyl through Galaxy has some disadvantages. Notably, Galaxy is more complex and so more cumbersome to install than a simpler command-line based pipeline. To address this we have made SNVPhyl available as simple to install virtual machine and Docker images, although these options cannot be straightforwardly implemented in a high performance computing environment.

Several factors can influence the ability to accurately call SNVs when using a reference mapping approach (39). As well, there are aspects of the datasets—such as recombination and population diversity—that can influence the phylogenetic analysis of identified SNVs. To assist in selecting proper parameters for SNVPhyl and gauging performance on different datasets we have assessed SNVPhyl under a variety of situations: SNV calling accuracy with simulated data, recombination masking, and the ability to differentiate outbreak isolates from non-outbreak isolates under differing parameters and data qualities.

Our assessment of SNV calling accuracy shows that SNVPhyl can detect SNVs and produce a SNV alignment with high sensitivity and specificity (Table 2). Of the variants that went undetected by SNVPhyl a large proportion were due to the quality thresholds and masking procedures implemented by SNVPhyl to remove incorrectly called or problematic SNVs (e.g., SNVs in internal repeats on the reference genome). While these quality procedures generate many false negatives they also eliminate many false positive variants—a reduction of 51 to 0 false positives at a cost of an additional 457 false negatives in the simulated dataset. However, all detected variation across all genomes is recorded in a table produced by SNVPhyl and additional software is provided for more detailed analysis of these variants.

Phylogenetics assumes descent with modification, but recombination (horizontal gene transfer) violates this assumption and its presence can confound the resulting phylogeny leading to misinterpretations on the clonal relationship of isolates (40). Recombination detection software exists and can be used for the construction of phylogenetic trees based on vertically inherited information (18, 19, 41). However, these programs require the pre-construction of whole genome alignments and can only be run on a single machine, which limits their utility for routine application to large collections of WGS reads.

SNVPhyl implements a basic but rapid method for detection and masking of recombinant sites by searching for SNV-dense regions above a defined density in a sliding window. We evaluated SNVPhyl’s recombination-masking method in comparison to the Gubbins software package which was run on a previously generated whole-genome alignment (Table 3, Figure S1). We found that SNVPhyl removes the majority of recombinant SNVs (from 2,159 SNVs with no recombination masking to 6 SNVs when masking regions with 2 SNVs in a 1000 bp window). However, SNVPhyl also removes some non-recombinant SNVs (reduced from 142 SNVs with no masking to 125 SNVs with 2 SNVs in 1000 bp). Removal of a greater number of recombinant SNVs is possible by increasing the window size, but this removes additional non-recombinant SNVs and reduces the information available in the phylogenetic tree and so concordance with other recombination-masking procedures (based on K tree scores).

SNVPhyl’s method of detecting high-density SNV regions can be executed independently for each genome. Independent execution is easily distributed across multiple nodes within a compute cluster, enhancing the scalability over large datasets. However, SNVPhyl requires the SNV density to be set *a priori* and may not be appropriate for organisms with complex evolutionary dynamics or for genome sequences from organisms spanning a large phylogenetic distance. We suspect that the optimal parameters will vary based on the particular organism under study and we would caution against relying on default settings without further evaluation. SNVPhyl does not aim to be a rigorous recombination detection and removal software package. However, SNVPhyl provides output files recording all the SNVs detected, which can be used for further analysis if needed. In particular, additional tools are provided that can produce a whole genome alignment correctly formatted for input into software such as Gubbins for a thorough detection of recombination and construction of a phylogenetic tree from non-recombinant SNVs.

A proper interpretation of the produced phylogenetic trees and SNV distances for associating closely-related isolates requires knowledge of when to trust the results and when additional parameter or data adjustments are necessary. To assist in defining these criteria we evaluated the performance of SNVPhyl at clearly delineating different outbreak clades across four different scenarios (Table 4, Figure S2).

In both the first and second scenarios we examined the effect of sequencing coverage on identifying enough SNVs to properly differentiate outbreak isolates. In the first scenario, we adjusted the minimum base coverage required to call a SNV from 5X to 20X without any additional subsampling of reads. We found that SNVPhyl succeeded in differentiating outbreak isolates at coverages up to 15X, but at a minimum coverage of 20X SNVPhyl failed to differentiate the outbreak isolates due to removal of too many SNVs (from 317 SNVs to 165 SNVs). In the second scenario, we subsampled one of the isolates along the mean read coverage values from 30X to 10X while keeping the minimum base coverage parameter in SNVPhyl fixed at 10X. We found SNVPhyl succeeded in differentiating outbreak isolates at a mean coverage of 15X and above, but failed to differentiate outbreak isolates at a mean coverage of 10X due to removal of too many SNVs (reduced from 299 SNVs to 155 SNVs). Both cases show that a high base coverage threshold for variant calling relative to the mean coverage of the lowest sample leads to falsely identifying samples as being related due to removal of too many SNVs (20X minimum coverage / 30X lowest sample mean coverage for failure in the first scenario, and 10X minimum coverage / 10X lowest sample mean coverage for failure in the second scenario). However, a high minimum base coverage threshold or too little sequencing data can be detected by examining the percentage of the reference genome considered as part of the core genome by SNVPhyl. A low value can indicate either a poorly related reference genome, or that large portions of the genomes are removed from the analysis (drop from 95% to 54% in the first scenario and 92% to 47% in the second scenario). We would recommend searching for such low values in the percent core to gauge whether or not base coverage (or possibly reference genome selection) is an issue for the SNVPhyl results.

In the third scenario (Table 4, Figure S2.c) we adjusted the SNV abundance ratio among values from 0.25 to 0.9. We found that SNVPhyl successfully differentiated outbreak isolates above a ratio of 0.5, but at a ratio of 0.5 the maximum SNV distance between isolates within an outbreak exceeded our threshold of less than 5 SNVs. However, unlike the minimum base coverage value, the percent of the reference genome identified as the core genome remained the same (92%). We recommend keeping this setting fixed at a higher value, with the default set at 0.75.

In the fourth scenario we simulated contamination between two closely-related isolates from two different outbreaks by mixing reads at differing proportions (Table 4, Figure S2.d). Our findings indicate that SNVPhyl is able to handle low amounts of mixed sample contamination (up to 10%). A higher proportion of contaminated reads can lead to removal of SNVs due to not meeting quality thresholds (from 298 SNVs with 5% contamination to 260 SNVs at 20% contamination where failure occurred) and so incorrectly implying relatedness between samples. Similar to the third scenario, the percentage of the reference genome identified as the core genome remained fixed at 92%. While SNVPhyl is able to differentiate outbreak isolates at low levels of contamination SNVPhyl cannot be used to evaluate the degree of contamination. Thus, we would not recommend the straightforward application of SNVPhyl to contaminated datasets without further assessment of the degree of contamination, either through taxonomic identification software such as Kraken (42); or, for closely-related isolates, through inspection of the variant calling and read pileup information provided by SNVPhyl.

Our analysis suggests that great care must be taken to reduce sources of noise in genome-wide SNV analysis. Some of this noise relates to quality thresholds for calling high quality SNVs, of which a careful balance is required to eliminate false positives without removal of too many true variants. Other sources include aspects of the WGS datasets or organisms under study such as the presence of contamination or recombination. The studied cases highlight how SNVPhyl is able to produce accurate phylogenetic trees under a wide variety of data qualities, and demonstrate how to detect inaccurate trees using additional information generated by SNVPhyl.

## Conclusion

SNVPhyl provides an easy-to-use pipeline for processing whole genome sequence reads to identify SNVs and produce a phylogenetic tree. We have shown that SNVPhyl is capable of producing accurate results on even very closely related bacterial isolates under a wide variety of parameter settings and sequencing data qualities. SNVPhyl is distributed as a pipeline within Galaxy and is integrated within the IRIDA platform, providing a “push button” system for generating whole genome phylogenies within a larger WGS data management and genomic epidemiology system designed for use in clinical, public health, and food regulatory environments.

## Authors contributions

A.P., N.K., C.N., A.R., E.T., R.B., W.H., F.B., M.G., and G.V.D. wrote the manuscript. A.P., L.K., A.R., and G.V.D. contributed to the initial software design while A.P., P.M., C.S., N.K., A.R., K.W., R.Z., and G.V.D. extended the design to a fully automated pipeline. A.P., P.M., C.S., and G.V.D. wrote the software with L.K., J.C., Mari. I., and Mark I. providing additional scripts or wrapping Galaxy tools. A.P., C.S., A.R., K.W., and G.V.D. designed the validation experiments. A.P. and C.S. performed the validation experiments. A.P. and G.V.D. interpreted the results.

## Acknowledgments

The authors would like to acknowledge Cheryl Tarr for initial inspiration and contribution to the design of SNVPhyl as well as Lauren Sluskey and Brian Yeo for their contributions during development of the pipeline. The authors would also like to acknowledge the Galaxy team and Galaxy community for their rapid response to issues and feature requests during the development of SNVPhyl as well as the integration of some bioinformatics tools within Galaxy that are used by SNVPhyl.

## Supplementary Materials

Supplementary materials are available in a separate file.

